# Cryo-EM sample preparation method for extremely low concentration liposomes

**DOI:** 10.1101/494997

**Authors:** Lige Tonggu, Liguo Wang

## Abstract

Liposomes are widely used as delivery systems in pharmaceutical, cosmetics and food industries, as well as a system for structural and functional study of membrane proteins. To accurately characterize liposomes, cryo-Electron Microscopy (cryo-EM) has been employed as it is the most precise and direct method to determine liposome lamellarity, size, shape and ultrastructure. However, its use is limited by the number of liposomes that can be trapped in the thin layer of ice that spans holes in the perforated carbon film on EM grids. We report a long-incubation method for increasing the density of liposomes in holes. By increasing the incubation time, high liposome density was achieved even with extremely dilute (in the nanomolar range) liposome solutions. This long-incubation method has been successfully employed to study the structure of an ion channel reconstituted into liposomes. This method will also be useful for preparing other biological macromolecules / assemblies for structural studies using cryo-EM.

## 1. Introduction

In the past four years, about 1,000 protein structures have been determined to better than 4-Å resolution in the EM databank using cryo-Electron Microscopy (cryo-EM). Among them, many are of detergent-solubilized membrane proteins (Fan et al., 2015; Hite et al., 2017; Hite et al., 2015; Liao et al., 2013; Tao et al., 2017; Yan et al., 2015). However, the use of detergent to solubilize membrane proteins always raises a question about whether or not the protein structure represents a biologically relevant state. As shown by both structural and functional studies, lipid membrane environment plays an essential role for the structural integrity and activity of membrane proteins (Gonen et al., 2005; Hilgemann, 2003; Hille et al., 2015; Lee, 2011; Long et al., 2007; Schmidt et al., 2006). One method is to use lipid nanodiscs, where the membrane protein resides in a small patch of lipid bilayer encircled by an amphipathic scaffolding protein (Bayburt et al., 2002). The nanodisc method has been employed to study the anthrax toxin pore at 22-Å resolution (Katayama et al., 2010), TRPV1 ion channel in complex with ligands at 3–4 Å resolution (Gao et al., 2016), and other membrane proteins (Autzen et al., 2018; Bayburt et al., 2002; Chen et al., 2017; Dang et al., 2017; Gao et al., 2016; Jackson et al., 2018; Katayama et al., 2010; McGoldrick et al., 2018; Roh et al., 2018; Srivastava et al., 2018; Taylor et al., 2017). Another method, called “random spherically constrained” (RSC) single-particle cryo-EM, where the membrane protein is reconstituted into liposomes, was developed and employed to study the large conductance voltage- and calcium-activated potassium (BK) channels reconstituted into liposomes at 17-Å resolution (Wang and Sigworth, 2009). Although both the nanodisc and RSC methods restore the lipid environment of membrane proteins, there is a major difference: the RSC method mimics the cell and provides an asymmetric environment (i.e. inside and outside conditions can be varied independently), whereas there is only one environment surrounding the membrane proteins in the nanodisc method. This is especially important for membrane proteins where an asymmetric environment is preferred (e.g. applying ligands to only one side of the membrane or applying transmembrane potential for voltage-gated ion channels or voltage-sensitive proteins, or applying pressures to mechanosensitive channels).

Liposomes are widely used as carrier systems in pharmaceutics (e.g. encapsulate therapeutic agents for cancer therapy (Børresen et al., 2018; Yuba, 2018; Zununi Vahed et al., 2017) and for neurological diseases (Vieira and Gamarra, 2016)), food industries (e.g. encapsulate bioactive food compounds to improve flavoring and nutritional properties or protect the food from spoilage and degradation (Shukla et al., 2017)) and cosmetics (e.g. as carriers of vitamins (Mozafari, 2005; Shashi et al., 2012). Liposomes have been proven to be beneficial for stabilizing encapsulated agents, overcoming obstacles to cellular uptake, and improving delivery efficiency of compounds to targeted sites *in vivo*. It is critical to accurately characterize liposomes and drug-liposome interactions as biophysical properties of liposomes are known to influence biological activity, biodistribution, and toxicity. Among the available techniques (Dynamic Light Scattering (DLS), Size Exclusion Chromatography (SEC), Atomic Force Microscopy (AFM), and cryo-EM), cryo-EM is the most precise and direct method to determine liposome lamellarity, size, shape and ultrastructure, which may reveal clues to mechanism of action toward the clinical endpoints of efficacy and toxicity(Aissaoui et al., 2011; Al-Ahmady et al., 2016; Baxa, 2018; Bonnaud et al., 2013; Crawford et al., 2011; De Carlo et al., 2004; Lepault et al., 1985; Uhl et al., 2017; Uhl et al., 2016; Zhang et al., 2013).

A major barrier for characterizing liposomes using cryo-EM is the difficulty of achieving high number of liposomes in the holes in the carbon film coated on TEM grids (Haghiralsadat et al., 2017; Park et al., 2014; Wang et al., 2008). Previous methods used very high concentration of liposome (Frederik and Hubert, 2005) or continuous carbon film coated grids (Jensen et al., 2016) or 2D crystal coated grids (Wang et al., 2008) or graphene coated grids (Palovcak et al., 2018). These methods either require high concentration of liposomes which is not trivial to achieve when membrane proteins are reconstituted into liposomes, or result in higher background noise due to the usage of extra layer of substrate. Inspired by the work by Snijder et al (Snijder et al., 2017) where multiple rounds of application and blotting (termed as multiple-application method) were used to reduce the required sample concentration, we present a long-incubation method to increases liposome density in holes even with extremely dilute liposome solutions. Based on systematic tests, the mechanism of the multiple-application and the long-incubation methods was discovered: the saturation of carbon regions by liposomes (i.e. monolayer coverage) only determines whether or not liposomes go into holes, but the accumulation of liposomes (i.e. multilayer adsorption) in carbon regions determines how many liposomes go into holes.

## 2. Methods

### 2.1 Liposome preparation

Liposomes were prepared in two ways: gel-filtration and extrusion (the resulting liposomes are termed as Liposome G and E, respectively). In gel-filtration method, 1-palmltoyl-2-oleoyl-sn-glycero-3-phophocoline (POPC) from Avanti (Alabaster, Alabama) was dissolved in buffer A (150 mM KCl, 20 mM HEPES, pH 7.3) in the presence of n-Decyl-β-D-maltopyranoside (DM) from Anatrace (Maumee, Ohio) with a lipid:detergent molar ratio of 1:3. Then appropriate 0.25 ml of the lipid-detergent mixture containing 0.4 mM lipid and 4 mM DM in buffer A (150 mM KCl, 20 mM HEPES, pH 7.3) was slowly loaded upon the top of a Sephadex G-50 (Sigma Aldrich, St. Louis, MO) column (hand packed in a 10 mm I.D x 300 mm height Econo column, 24 ml volume), which had been pre-equilibrated with running buffer A. The flow was driven by gravity at a speed of 0.2–0.3 ml/min, and elution fractions were monitored by ÄKTA purifier system (GE Healthcare, Chicago, IL) and collected by Frac 920 (GE Healthcare, Chicago, IL). In extrusion method, POPC lipid dissolved in chloroform was dried under nitrogen, and re-hydrated in buffer A. The lipid suspension was frozen and thawed 10 times, and extruded through an 50-nm polycarbonate membrane filter (Whatman) using a Lipex™ extruder (Northern Lipids Inc.) (Mayer et al., 1986). Then the lipid concentration was adjusted to 2 mM by concentration. All operations were carried out at Room Temperature.

### 2.2 Expression, purification and reconstitution of human BK proteins

The BK proteins were expressed and purified as previously described (Wang and Sigworth, 2009). Briefly, full-length human Slo protein (GI:507922) carrying an N-terminal FLAG tag was stably expressed in HEK293 cells. The BK protein was purified using an anti-FLAG affinity column (Sigma Aldrich), where the detergent dodecyl maltoside (DDM) (Anatrace) was exchanged with decyl maltoside (DM) (Anatrace) before elution with FLAG peptide (Sigma Aldrich). Superose6 was used to further purify tetrameric BK.

To reconstitute the BK protein for structural study, BK peak after Superose6 was mixed with DM-solubilized POPC (1-palmitoyl-2-oleoyl-sn-glycero-3-phosphocholine) lipid (Avanti Polar Lipids, Inc.) (DM: POPC = 3:1) giving a final protein-to-lipid molar ratio of 1:1,000. Gel filtration on a hand-packed 24ml (10mm I.D and 300mm height) was used to remove detergent with running buffer containing 20 mM HEPES, pH 7.3, 150 mM KCl, 2 mM EDTA. To reconstitute the BK for functional assay, the purified BK were reconstituted into liposomes as previously described (Li et al., 2015). Briefly, the POPC and POPG (1-palmitoyl-2-oleoyl-sn-glycero-3-phospho-(1'-rac-glycerol)) lipid mixture (3:1 molar ratio) in chloroform was dried under nitrogen for 30 min, and rehydrated and sonicated in Fisher Scientific FS20H Ultrasonic Cleaner (Fisher Scientific) in buffer A (20 mM HEPES, pH 7.3, 150 mM KCl) to a final lipid concentration of 10 mM. The resulting solution was mixed with detergent DM (lipid : DM = 1 : 1) for 30 min. Then the purified BK channels were added to a final protein concentration of 0.05 mg/ml. The detergent was removed by dialysis in 12–14 KD cut-off dialysis membrane (Spectrum Labs) against the buffer A at 4 °C for 3 days (dialysis solution was changed every day). Empty liposomes were prepared in the same manner without the addition of protein prior to dialysis.

### 2.3 Cryo-EM sample preparation, image collection and analysis

Three protocols were used to prepare cryo-EM samples: (A) on continuous carbon film coated grids: 2 μl of liposome solution was applied onto a glow-discharged continuous carbon coated TEM grid, blotted and fast frozen at 22 °C and 100% relative humidity in an FEI Vitrobot Mark IV (FEI) as for normal cryo-sample preparation. (B) multiple-application method: 2 μl of liposome solution was applied onto a glow-discharged holey carbon grid (CFlat R2/2, EMS) and blotted briefly from the edge. This procedure was repeated to the desired number of rounds of sample application (Snijder et al., 2017). After the last round of sample application, the sample was blotted and fast frozen at 22 °C and 100% relative humidity in an FEI Vitrobot Mark IV (FEI) as for normal cryo-sample preparation. (C) long-incubation method: 2 μl of liposome solution was applied onto a glow-discharged holey carbon grid (CFlat R2/2, EMS), and incubated for 0.25-10 min at 22 °C and 100% relative humidity in an FEI Vitrobot Mark IV (FEI). After a brief blotting from the side of the grid, another 2 μl of solution with or without liposomes was applied to the grid. After 10-20s, the grid was blotted and frozen as for normal cryo-sample preparation. Samples were imaged on an FEI G2 F20 (FEI, Hillsboro, OR) equipped with Eagle 4k camera (FEI, Hillsboro, OR) or an FEI T12 (FEI, Hillsboro, OR) equipped with a Gatan US4000 CCD camera (Gatan, Pleasanton, CA). The dose for each exposure was about 2,000 e^-^/nm^2^. Images were taken at 67,000 (T12) or 50,000 (F20) magnification and 2.0-3.3 μm underfocus. Liposomes in each cryo-EM image were identified automatically using a home-made Matlab program as previously described (Wang et al., 2006). Then the size distribution and mean size was analyzed using Matlab.

The liposome concentration is estimated as outlined below. The average number of lipid molecules per liposome is:

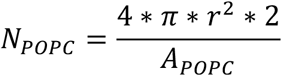

where *r* is the radius of a liposome (from the center of the liposome to the center of the bilayer), *A_POPC_* is the area per lipid molecule (68 Å^2^) (Kučerka et al., 2006). The liposome concentration is:

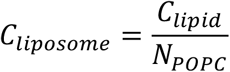

where *C_lipid_* is the lipid concentration.

## 3. Results

### 3.1 Multiple-application method

Liposomes were prepared using both extrusion (Liposome E) and gel filtration (Liposome G) methods. Both methods produced unilamellar liposomes as shown in Fig. 1. The mean average sizes are 270 and 90 Å in radius for Liposome E and G, respectively. Cryo-samples were prepared using multiple-application method as previously described (Snijder et al., 2017). With one round of sample application (i.e. normal cryo-sample preparation method), there were few liposomes in holes (Fig. 2A&D) for both Liposome E and G. After 6 rounds of sample application, more liposomes went into holes for Liposome G (Fig. 2 A-C). However, even after 6 rounds of sample application, only a few liposomes went into the holes for Liposome E (Fig. 2 D-F). This is due to the lower liposome concentration of Liposome E (although Liposome E and G have the same lipid concentration, the liposome concentration of Liposome E is about 9 times lower than that of Liposome G (66 nM vs. 590 nM)).

**Figure 1.**
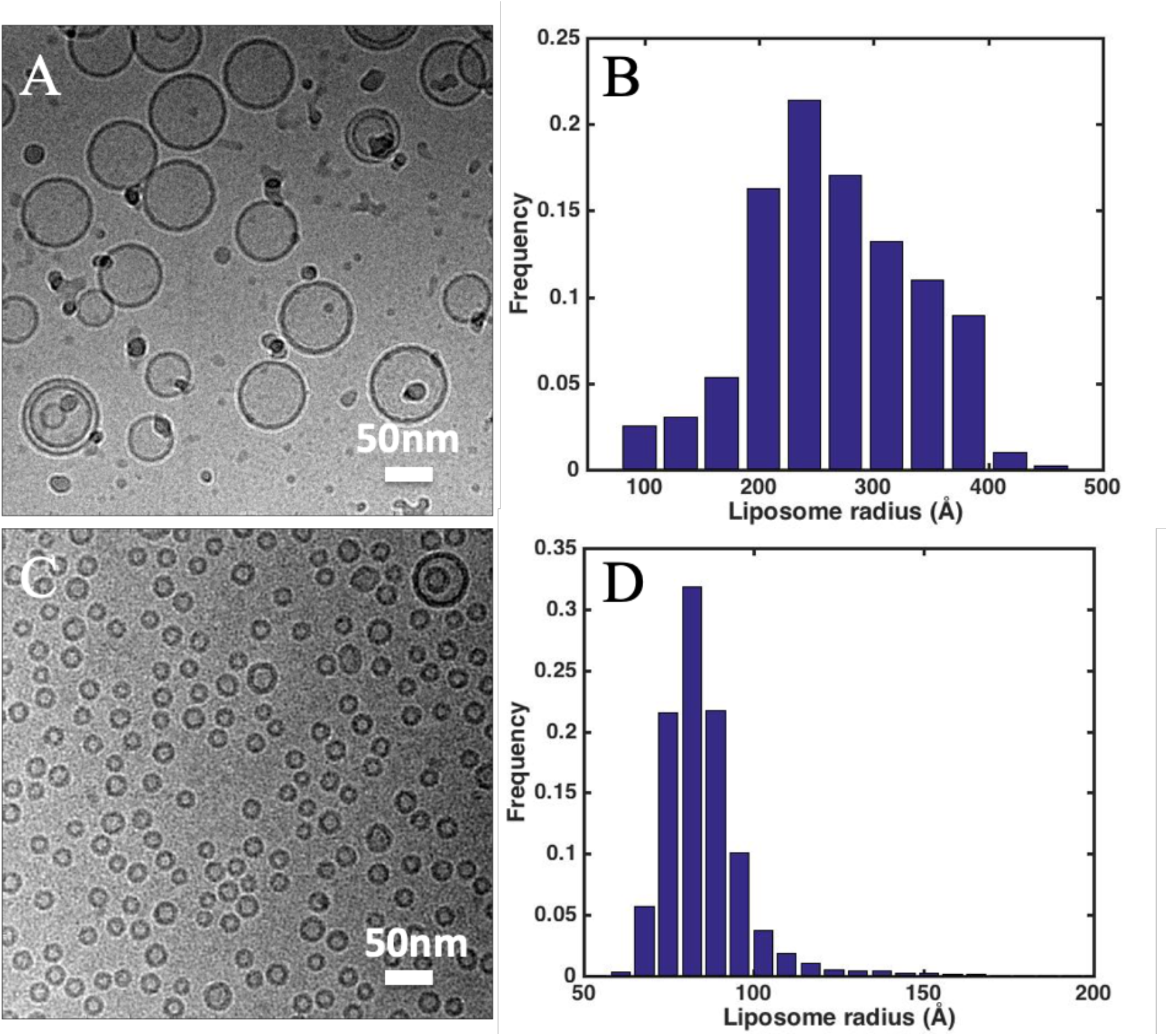
Cryo-EM images and size distribution of (A-B) extruded liposomes (Liposome E) and (C-D) gel-filtration liposomes (Liposome G) applied to TEM grids coated with continuous carbon film. The mean radius of Liposome E and G are 270 and 90 Å, respectively 90.

**Figure 2.**
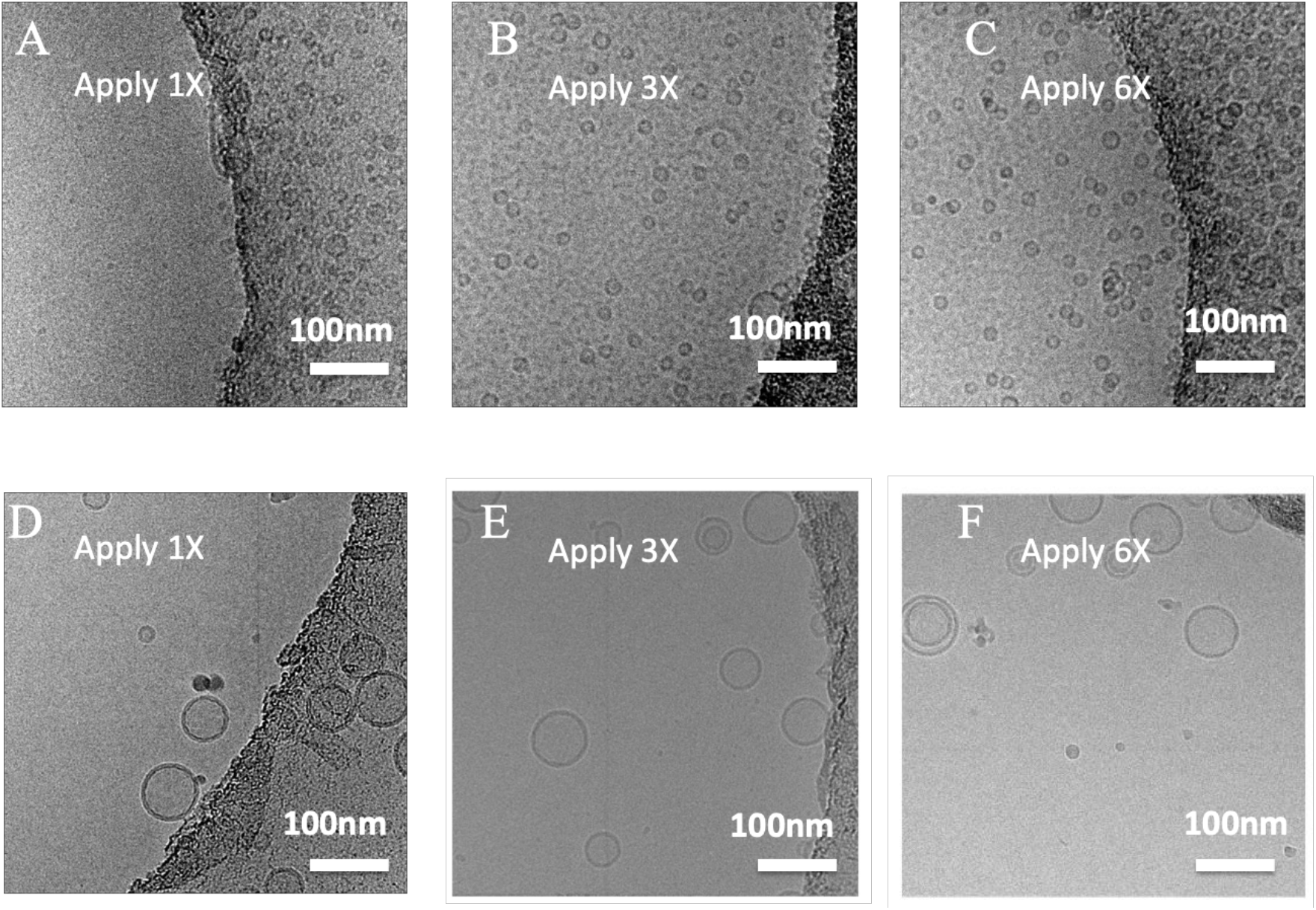
cryo-EM images of liposomes by multiple-application method. (A-C) Liposome G was applied once, three times and six times. (D-F) Liposome E was applied once, three times and six times. Scale bar: 100 nm.

### 3.2 Long-incubation method

Although multiple-application of samples increased the liposome density in holes to some extent as seen in Fig. 2C for Liposome G, it did not work for Liposome E which has a lower liposome concentration. To reduce the required liposome concentration and the amount of sample to be used (i.e. one-time sample application *vs* six-time sample applications), a long-incubation method was developed. As shown in Fig. 3A, with a one-minute incubation, the liposome density in holes was comparable to that prepared by multiple-application method (Liposome G was applied six times) (Fig. 2C). When the incubation time increased to 10 min, the liposome density increased by a factor of three (Fig. 2C). Similar phenomena were observed for Liposome E. With a 10-minute incubation, many liposomes went into holes for Liposome E (Fig. 3D). The long-incubation method reduced the requirement for both concentration and amount of liposome solutions.

**Figure 3.**
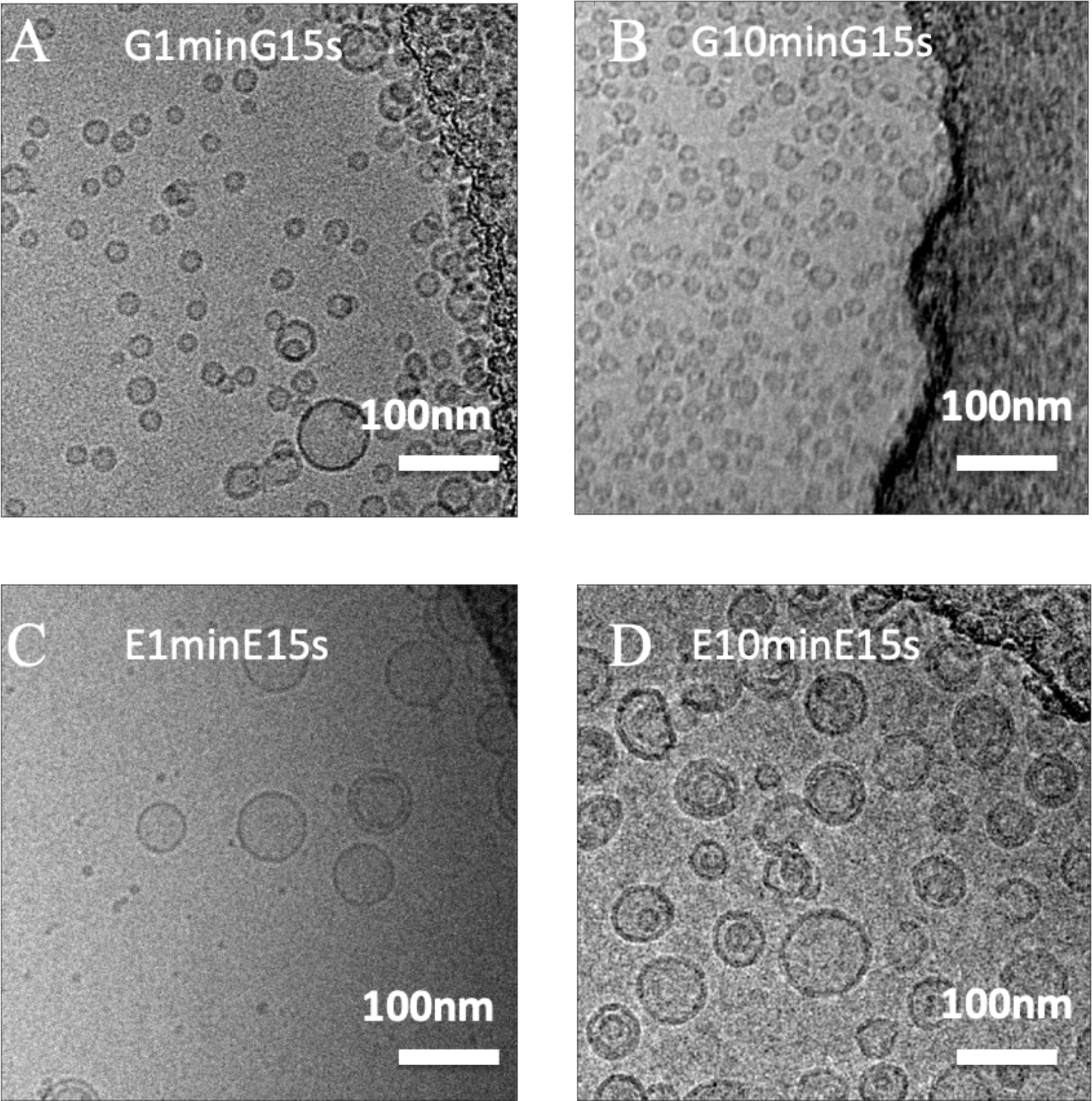
cryo-EM images of liposomes prepared using the long-incubation method. (A-B) Liposome G was incubated for one minute (G1minG15s) and 10 minutes (G10minG15s) before blotting. Then Liposome E was applied again, incubated for 15 s, blotted and fast frozen. (C-D) cryo-EM samples of Liposome E were prepared using the same method as in A and B.

### 3.3 Long-incubation followed by an application of different liposome solutions

As shown in Fig 3, long incubation increased liposome density in holes for both Liposome G and E. To track whether the increased liposome density came from the first round or the second round of sample application, Liposome E was added in the first round of sample application and Liposome G was added in the second round of sample application. As shown in Fig 4A (one-minute incubation), the carbon region of the grid was not fully covered by liposomes and few liposomes populated in holes. When the incubation time increased to five minutes, the carbon region was saturated with liposomes (fully covered), and some liposomes went into holes (Fig. 4 B). With a 10-minute incubation, much more liposomes went into the holes (Fig. 4C). This suggested that the accumulation of liposomes (i.e. multilayer adsorption) in carbon regions is critical to have liposomes populated in holes.

**Figure 4.**
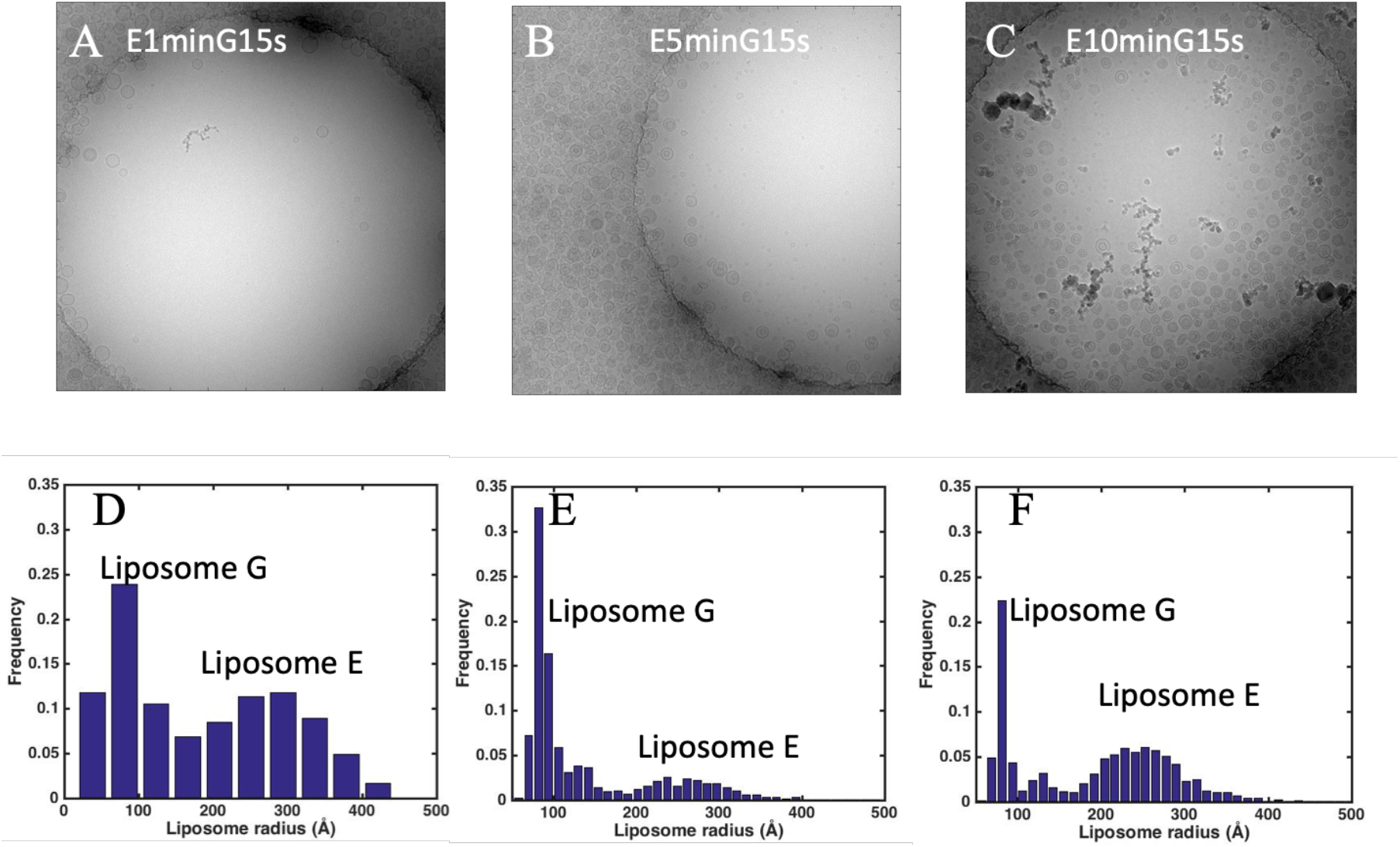
Contributions of the first and second application of samples on liposome density in holes on a holey-carbon grid. (A-C) representative cryo-EM image prepared with one-, five-, 10-minute incubation of Liposome E followed by one application of Liposome G. (D-F) liposome sizes calculated from cryo-EM images prepared as shown in A-C. Holes are 2 μm in diameter. The mean sizes of Liposome E and G are 270 and 90 Å, respectively. Please note the carbon region in A was not fully covered by liposomes, while the carbon region in B and C were fully covered by liposomes.

As shown in Fig. 4 D-F, two liposome populations were observed: Liposome E with a mean radius of 270 Å and Liposome G with a mean radius of 90 Å. The liposome coverage due to Liposome E increased from 2.5% to 6% and 18% when the incubation time increased from one minute to five and ten minutes (Fig. 6 B). However, the liposome coverage due to Liposome G was the same (~2%) with five-minute and 10-minute incubation of Liposome E as the incubation time for Liposome G was the same. The reason for a lower coverage of Liposome G in Fig. 4A lies in the fact that the carbon region was not saturated by Liposome E with the one-minute incubation. Newly added Liposome G preferably went to carbon regions.

### 3.4 Long-incubation followed by an application of liposome-free solution

As suggested by the results shown in previous sections, a long incubation time after the first application of liposome solution was critical to increase the liposome density observed using cryo-EM. The long incubation not only saturated the carbon regions (i.e. monolayer coverage), but also make liposomes accumulate/pile up in carbon regions (i.e. multiplayer adsorption). The second application of sample flushed the accumulated liposomes from carbon regions into holes. To confirm this hypothesis, liposome-free buffer A was used in the second application of sample. As shown in Fig. 5, even with the liposome-free buffer A, many liposomes were observed holes. In combination with the results shown in Fig. 4 D-F and Fig. 6 B-C, it is clear that saturation of carbon regions and accumulation of liposomes in carbon regions are both due to the first application of liposome sample.

**Figure 5.**
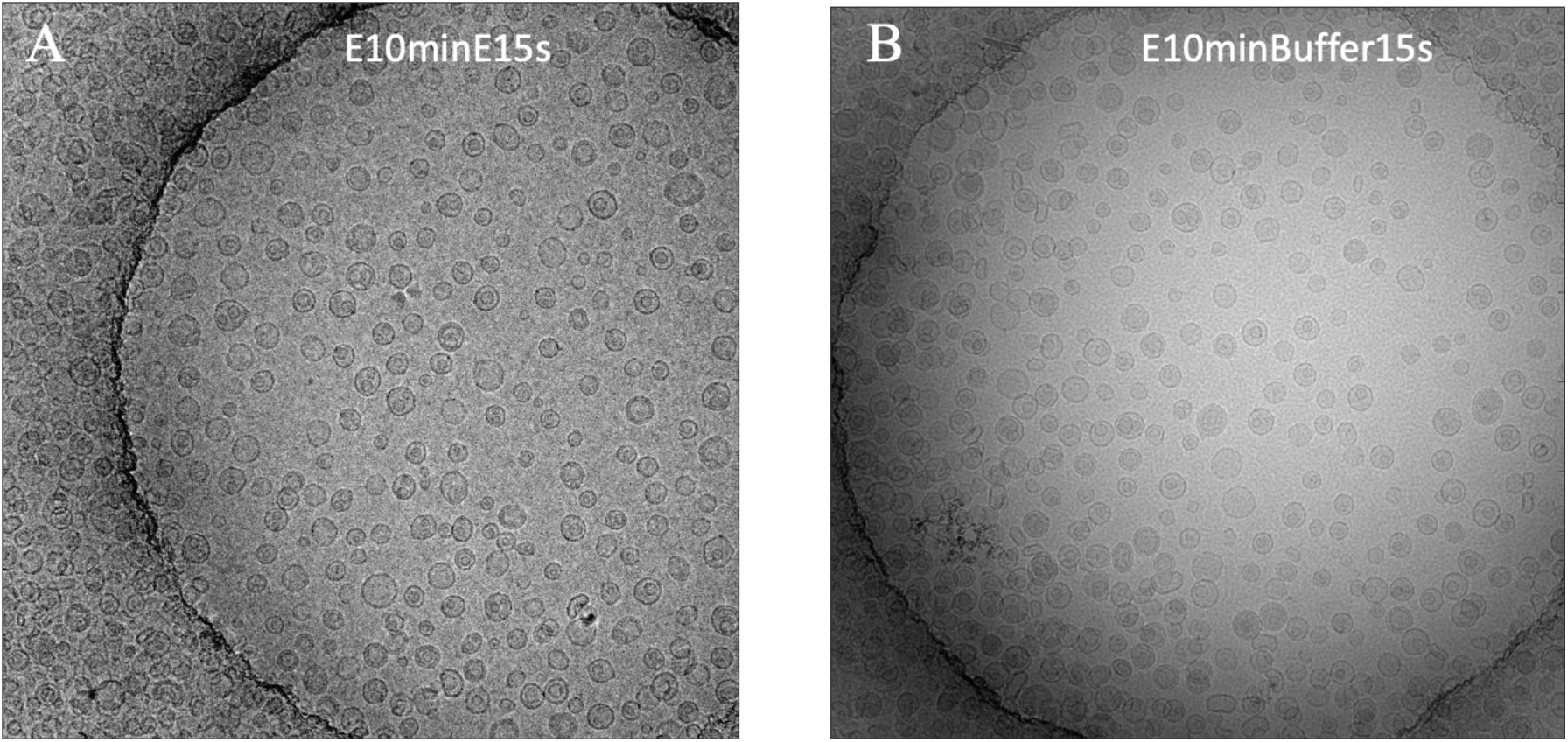
Cryo-EM images of Liposome E with 10-minute incubation followed by an application of (A) Liposome E or (B) liposome-free buffer A. Holes are 2 μm in diameter.

**Figure 6.**
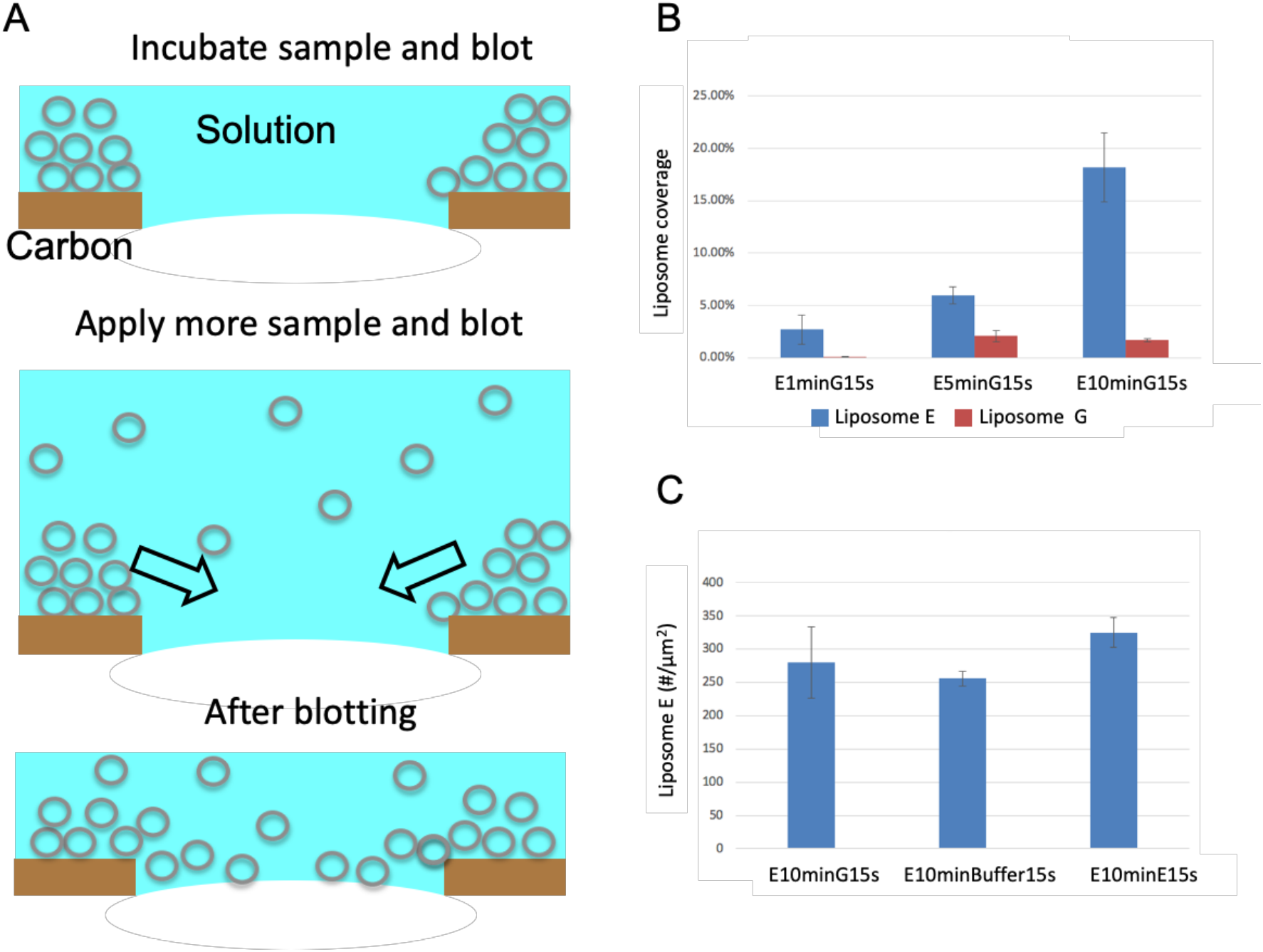
Mechanism of the long-incubation method. (A) A cartoon showing the long-incubation method. (B) Liposome coverage with different incubation times. (C) Liposome density when different solutions (Liposome G, liposome-free buffer A, and Liposome E) were used in the second sample application.

## 4. Discussion

The liposome density observed in holes on the EM grid depends on the bulk liposome concentration, the interaction between liposomes and the carbon film, and the blotting process. Using the same sample and the same standard cryo-sample preparation method, there are at least 20 times more liposomes on carbon film (Fig. 1 A&C) than in holes (Fig. 2 A&D). However, the carbon film increases the background noise, thus decreases the signal-to-noise ratio (SNR) of the cryo-EM images, which is inferior for high-resolution studies (e.g. interaction between drugs and liposome, the structure of reconstituted membrane proteins). One remedy is to decrease the carbon film thickness: coating an ultra-thin carbon film over a holey carbon grid (Jensen et al., 2016). Another technique is to use multiple-application method (Snijder et al., 2017). The multiple-application method has been successfully employed to obtain high densities of biological macromolecules in holes with dilute samples (e.g. in micromolar range). However, this method did not work for Liposome E (e.g. in nanomolar range) as shown in Fig. 2 D-F.

As shown by the number of liposomes in carbon region (Fig. 1A&C) and in holes (Fig. 2 A&D), liposomes prefer carbon film over holes. As the incubation time increases from 15 s to 10 minutes, more and more liposomes accumulated/piled in carbon regions: there is still unoccupied area in carbon regions with one-minute incubation (top right corner of Fig. 4A), while liposomes overlapped with each other with five-min incubation (left portion of Fig. 4B). As the majorities of liposomes are unilamellar (Fig. 1A&B), the overlapping of liposomes indicates that liposomes are piled up on carbon film as illustrated in Fig. 6A. After the first blotting, a thin layer of aqueous solution is left on the grid. With a second application of sample and blotting, the accumulated/piled liposomes in carbon regions are flushed into holes. The liposomes in holes is a combination of the accumulated liposomes from the first sample application and the liposomes from the second sample application. As shown in Fig. 6B, the liposome coverage due to the first application of sample (Liposome E) increased with increased incubation times. The liposome coverage due to the second application of sample (Liposome G) is constant as it is determined by the bulk liposome concentration of the applied liposome solution. However, if the incubation time is not long enough and the carbon region is not saturated (i.e. not fully covered), the liposomes from the second application of sample will preferably go to carbon regions to saturate the carbon film. Thus, the liposome coverage due to the second application of sample (Liposome G) is much lower with one-minute incubation than that with five-minute and 10-minute incubation of the first application of sample (Liposome E). To distinguish whether or not the accumulation of liposomes from the first application of sample determined the number of liposomes entering holes, a liposome-free buffer A was used in the second application of sample. A slightly lower liposome density was observed (Fig. 5B). As shown in Fig. 6C, the liposome density due to the first application of sample is lower (256±11) when liposome-free buffer A was used in the second sample application than when a different liposome solution was used in the second sample application (280±53). When the same liposome solution was used in the second sample application, a higher liposome density was observed (325±22). In short, the long-incubation not only saturates the carbon film, but also increases the number of accumulated liposomes in carbon regions. The accumulation of liposomes is the real step which increases the liposome density in holes. This is also the reason why the multiple-application method worked with dilute samples. The multiple application of dilute liposome solutions will saturate carbon regions (i.e. full monolayer coverage), but there is not enough time for liposomes to accumulate/pile in carbon regions. Thus a long-incubation time is necessary to saturate carbon films and accumulate liposomes in carbon regions, which will be flushed into holes later. Based on the similar effect of sample application numbers on liposome density and biological macromolecule/assembly density in holes and the similarity of lipids and proteins (i.e. all are biological samples), we believe this long-incubation method also work for other biological samples including proteins and viruses.

It’s worth noting that although the Vitrobot was set up at 100% humidity, opening and closing the chamber for tweezers movement may cause the humidity drop. When the relative humidity is less than 100%, there will be net water transport from the specimen to the environment, and the liposome will dehydrate. For an incubation time over five minutes, osmotic collapse of spherical liposomes into ‘vase’-like structures can be observed. One easy fix for this osmotic imbalance is to use a hypotonic solution in the second application of sample, which will swell the dehydrated liposomes (Wang and Sigworth, 2009). This long-incubation method has been successfully employed to study the structures of an ion channel reconstituted into liposomes. As shown in Fig. 7, a good distribution of proteoliposomes in holes was achieved. The protein particles were easily identified in the cryo-EM images before and after liposome subtraction.

**Figure 7.**
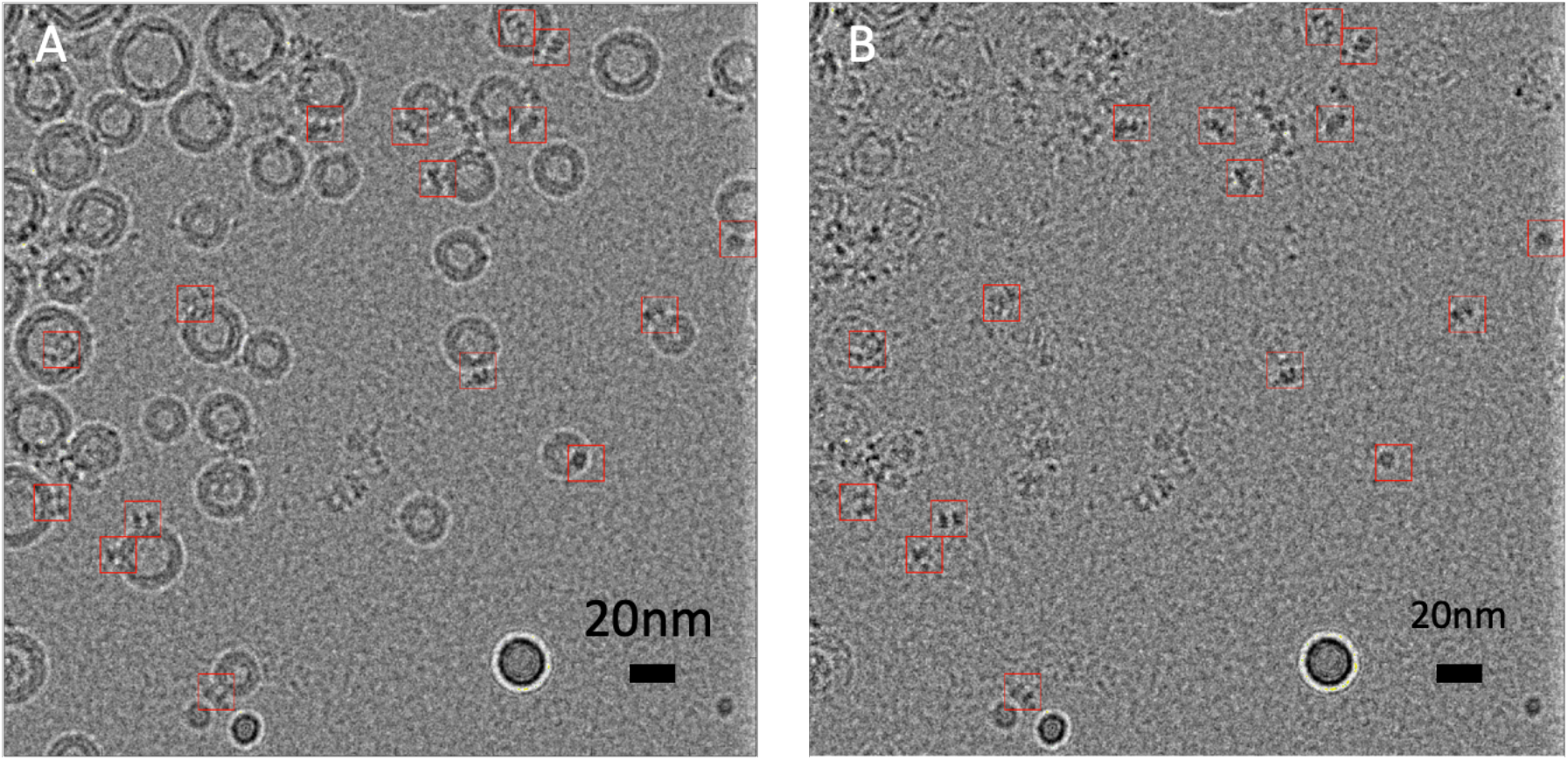
Liposome with reconstituted human BK channels prepared using long-incubation method. (A) A representative cryo-EM image of BK proteoliposomes. (B) Liposomes are subtracted from the image shown A. BK particles were marked with red boxes (15 nm).

## 5. Conclusion

Long-incubation method was developed to achieve desired liposome densities in cryo-EM images of liposomes even at extremely low liposome concentrations (e.g. in nanomolar range). It was employed to visualize liposomes produced by two commonly used methods (gel filtration and extrusion) and liposomes with reconstituted membrane proteins for high-resolution structural studies. We discovered that the saturation of carbon regions (i.e. monolayer coverage) determines whether liposomes go into holes, while the accumulation of liposomes in carbon regions (i.e. multilayer adsorption) determines how many liposomes go into holes. The long-incubation method is also applicable to prepare cryo-samples of other biological samples including proteins and nucleic acids.

## ACKNOWLEDGMENTS

This work was supported by the National Institutes of Health [grant number: R01GM096458].

## AUTHOR CONTRIBUTIONS

Conceptualization, L.W.; Methodology, T.L. and L.W.; Writing, T.L. and L.W.; Editing, L. T. and L.W.; Project Administration, L.W.; Funding Acquisition, L.W.

## DECLARATION OF INTERESTS

The authors declare no competing interests.

